# Elucidating the active interaction mechanism of phytochemicals withanolide and withanoside derivatives with human serum albumin

**DOI:** 10.1101/352575

**Authors:** Shreya Dubey, Monika Kallubai, Arijit Sarkar, Rajagopal Subramanyam

**Affiliations:** Department of Plant Science, School of Life Sciences, University of Hyderabad, Gachibowli, Telangana-500046 India

**Keywords:** *Withania somnifera*, Human serum albumin, Fluorescence spectroscopy, Molecular dynamics

## Abstract

*Withania somnifera* (Ashwagandha) is an efficient plant known in Ayurveda and Chinese medicine since ancient times, whose extracts are consumed orally as food supplement or as a health tonic owing to its several restorative properties for various CNS disorders, inflammation, tumour, stress and rheumatism. In this study, we have analyzed the binding interaction of four derivatives of *Withania somnifera* (Withanolide A, Withanolide B, Withanoside IV and Withanoside V) with HSA because of their important pharmacological properties. To unravel the binding between derivatives of *Withania somnifera* and HSA, fluorescence spectroscopy was used. Binding studies were further studied by molecular docking and dynamics studies and results confirmed greater stability upon binding of derivatives with HSA. Circular dichroism data illustrated change in the secondary structure of protein upon interaction with these derivatives, particularly the helical structure was increased and β-sheets and random coils were decreased. Furthermore, morphological and topological changes were observed using AFM and TEM upon binding of ligands with HSA indicating that HSA-withnoside/withanolide complexes were formed. All the results cumulatively demonstrate strong binding of withanosides and withanolides derivatives with serum albumin, which should further be explored to study the pharmacokinetics and pharmacodynamics of these *derivatives*.

## Introduction

Withanolides exits as secondary metabolites; structurally it consists of a steroidal backbone bound to a lactone or one of its derivatives generated from oxidation of steroids [1]. These compounds can be isolated from *Withania somnifera* (Indian Ginseng; family: *Solanacea*), also known as Ashwagandha, it is found to be loaded with numerous medicinal properties and useful in counteracting various illness like dehydration, muscle tension, memory loss and also restoring general health and vitality. This plant is known to be found globally, with more occurrences in xeric and drier regions of tropical and subtropical areas [2,3]. We have selected four derivatives of this plant, which are withanolide A (C_28_H_38_O_6_-470.6Da), withanolide B (C_28_H_38_O_5_-454.6Da), withanoside IV (C_40_H_62_O_15_-782.9Da) and withanoside V (C_40_H_62_O_14_-766.9Da) for our study (Scheme 1). The withanolides and withanosides are distinctive with their C-28 and C-40 steroidal lactones, with six-membered lactone ring formed by oxidization of the C-22 and C-28 on ergostane backbone. Although structurally similar, withanolide A differs from withanolide B with the presence of an extra hydroxyl group on C-20 atom, whereas withanoside IV differs from withanoside V with the presence of an extra hydroxyl group on C-27 atom The derivatives of *Withania somnifera* have been used since time immemorial in Ayurveda and ancient system of medicine for its pharmacological activities like chemopreventive, anti-inflammatory, anti-arthritic, and angiogenesis activity [4], and they are being used as a rasayana to promote immunity,health and longevity which acts as a adaptogen and immunomodulator. These compounds are also known to exhibit nootropic effect and are effective against several neurodegenerative diseases including Alzheimer’s and Parkinson’s disease [5], hence Ashwagandha derivatives have become very popular oral supplement globally because of its strong remedial properties. Owing to the remedial properties of withanolides and withanosides, their affinity to bind with human serum albumin (HSA) was carried out, which might play crucial role in its pharmacokinetics and pharmacodynamics.

**Scheme 1:**
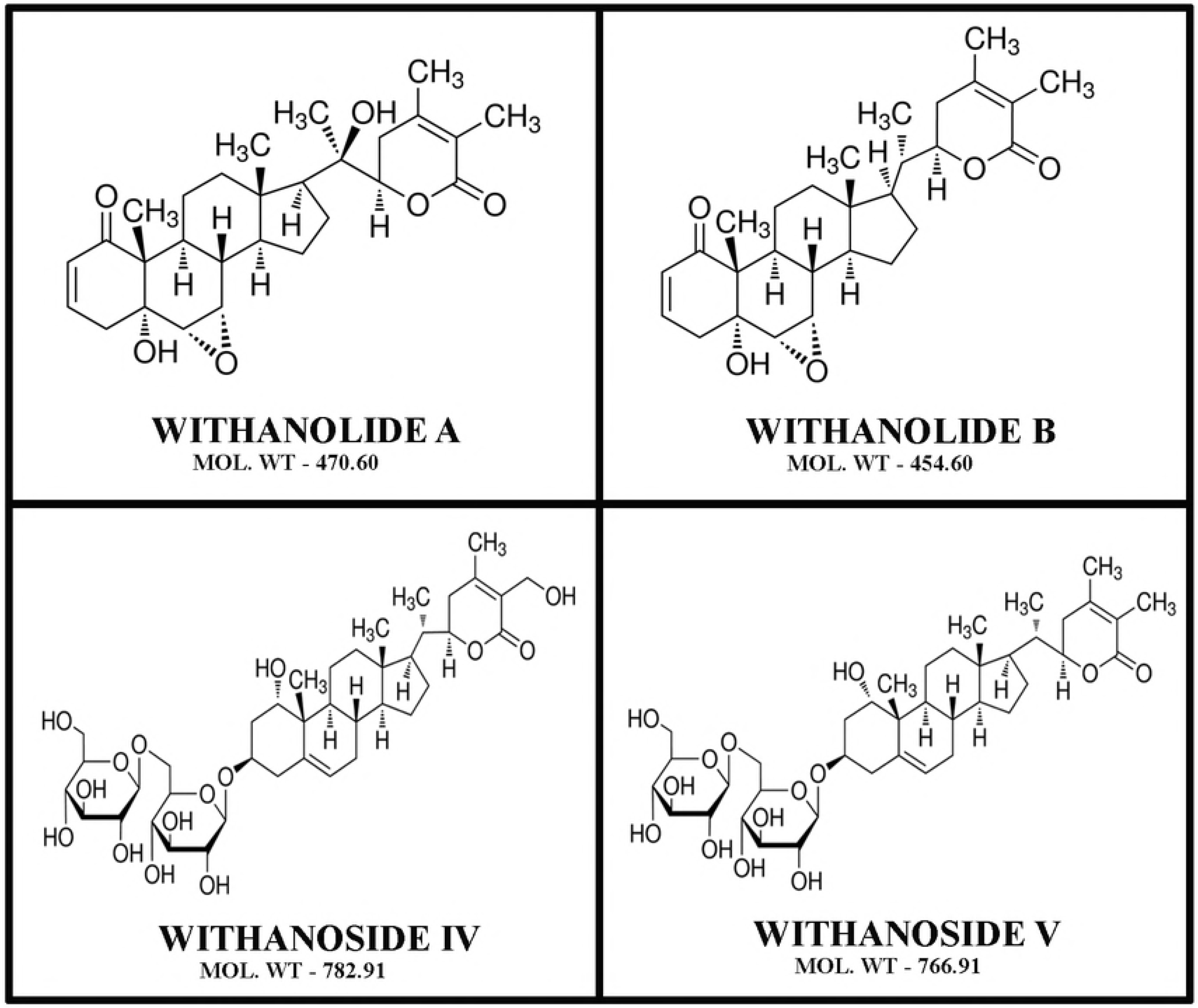
The structure of withanolide A (C_28_H_38_O_6_-470.6Da), withanolide B (C_28_H_38_O_5_-454.6Da), withanoside IV (C_40_H_62_O_15_-782.9Da) and withanoside V (C_40_H_62_O_14_-766.9Da). (E) HSA+withanolide V.

Albuminis a major circulating protein in blood plasma, comprising approximately 60% of the blood proteins [6,7] is a non-glycosylated protein of 66.5KDa, and 585 amino acids having 67% α-helix in its native conformation [8]. It has three domains I, II and III; each having two sub-domains A and B, which are connected via random coils. It has two important drug binding sites known as Sudlow's site II and I [9,10] which are domain IIIA, IIA and also domain IB noticed as one of drug binding site [11]. It is well established that HSA has 35 residues of Cys and forms 17 disulfide bonds that contributes to the stability of the protein. The single 34^th^residue of Cys imparts the protein its anti-oxidant property [12]. This negatively acute phase protein exerts pharmacological property because of its long half-life of 19 days, and its ability to bind with several exogenous as well as endogenous compounds [13]. The interaction between the drugs and plasma protein influences the absorption, bio-distribution, metabolism and excretion of the drug. Our group has earlier reported various therapeutic phytochemicals like lupeol, β-sitosterol, asiatic acid, trimethoxyflavone, and its derivative binds strongly to serum albumin [14–17].

Since withanolide is having several kind of biological importance and we presume that other four derivatives of withanolide A (C28H38O6), withanolide B (C_28_H_38_O_5_), withanoside IV (C_40_H_62_O_15_) and withanoside V (C_40_H_62_O_14_) also may have good therapeutic role. So far the role of these molecules in biological samples including *in vitro* or *in vivo* have not been explored. Since, human serum albumin is an important transport protein, we have chosen this protein to study the pharmocokinetics importance. Through our current study, we would decipher the binding mechanism, stability and conformation of these compounds with HSA.

## Materials and methods

### Preparation of stock solutions

Fat free HSA was procured from Sigma Aldrich and working concentration of 100μM was made by dissolving in 100mM PBS of pH 7.4 to maintain the physiological condition. Withanolides A, B and withanosides IV, V which are extracts of extracts of Indian Ginseng plant were procured from Natural Remedies Pvt. Ltd, Bengaluru, India with purity of ~99%. These compounds were first dissolved in dimethyl sulphoxide (DMSO) at 20 mM. This stock solution was further diluted to a concentration of 100μM working stock in PBS (pH 7.4) similar to our previously reported studies [18,19]. All other chemicals used were of analytical grade from Sigma Aldrich.

### Room temperature fluorescence spectra

Fluorescence spectroscopy is an astounding technique used to study the interaction between protein and ligands and calculation of binding constant and free energy. Concentration of HSA was kept constant at 1μM and Withanolides A&B and withanosides IV & V were titrated in an increasing concentration from 1μM to 9μM in 0.1mM PBS of pH 7.4. Perkin Elmer LS55 fluorescence spectrometer was used to record the spectra and incubation time for all the derivatives were kept constant for 5 mins. Emission spectra was recorded in the range of 300-500nm, with parameters set around like our previous studies [20]. Excitation wave length of HSA was kept at 285 nm and slit width 5 nm for excitation and emission spectra. Three independent experiments were carried out and each time spectra recorded was almost identical.

### Molecular displacement

For determining the exact binding site of the withanolides A&B and withanosides IV&V derivatives to HSA, we have studied the interaction of these derivatives with HSA in the presence of certain site specific markers (Lidocaine for domain I, Phenylbutazone for domain II, and Ibuprofen for domain III) [21–23]. The choices for the site specific individual markers were based upon extensive earlier binding site studies of different ligands with HSA. The concentration of HSA and the markers were kept constant at 1 μM, and remaining all parameters were same as fluorescence spectroscopy. Experiments were done in triplicates and based upon the binding energy values, the binding sites were determined.

### Circular dichroism measurements

For exploring the conformational change of the protein upon addition of a ligand, circular dichroism is referred to as the “Gold Standard”technique. Circular dichroism (CD) spectropolarimeter, Jasco-815 was used for the study and the temperature was kept constant at 25 °C. Spectra were recorded in a quartz cell with path length of 0.02cm. Data was recorded from 190-260nm with scan speed of 50nm/sec. The concentration of HSA was persistent at 1μM and the concentation of drugs 2, 4, 6 μM were added gradually similar to the previous studies [24]. CDNN 2.1 web based software was used to calculate the change in percentage of α-helix, β-sheet and random coils from its native conformation upon addition of withanolides and withanosides with increasing concentrations.

### Atomic force microscopy

Atomic Force Microscopy was used to visually detect any change in the topography of the protein molecule surface upon addition of the ligand. AFM experiments were done by NT-MDT solver scanning probe microscopy in semi contact mode using cantilever (0.3mm) with a force constant (5.5 −22.5N/m) and the typical imaging resonance frequency was 140 kHz. 3.6× 1.6×0.4mm was the size of the gold-coated silicon probes used; tip height was 14-16μm and radius of curvature 10 nm [25]. For sample preparation, 20 *μ*L of 2 *μ*M free HSA was spread on a glass slide and incubated for 10 minutes and then it was washed with 1ml de-ionized water to remove loosely bound molecules from the surface; and for HSA-ligand complexes, 20 *μ*L of a 10*μ*M withanolides and withanosides were first incubated with HSA and similarly spread on a glass slide, washed and dried under for 5 min. The dried samples were then imaged by AFM in non-contact mode.

### Transmission electron microscopy (TEM)

FE1 Tecnai G^2^S-Twin-200kv instrument was used to observe the change in the morphology of the protein upon addition of the ligand, in high resolution. 1μM HSA was used as the control and for HSA-ligand complexes, 1 μM HSA was incubated with 2 μM of withanolides (A and B) and withanosides (IV and V) individually. A drop of these samples was added on the carbon coated copper grids and excess sample was removed by absorption with filter paper. Before drying of the samples 2% uranyl acetate negative stain was added and final observation was made as described [19].

### Molecular docking

Molecular docking is a computation tool used to predict the predominant binding mode of the ligand to a protein and to locate geometrically and energetically the most stable conformer complex. Here, we have used Autodock 4.2.3 to generate the 50 docked conformers of the protein ligand complexes by using genetic lamarckian algorithm. The crystal structure of HSA (PDB ID: *1AO6*) was downloaded from the Brookhaven Protein Data Bank. Three-dimensional structure of withanolides and withanosides was built from 2D structure and geometry, and optimized using Discovery studio 3.5 software. Water molecules were removed, Kolmann charges and polar hydrogen atoms were added to the PDB structure of HSA before analysis. Grid size of 126 × 126 × 126 along X, Y and Z axis with 0.586 Å grid spacing was generated to carry out blind docking of withanolides and withanosides with HSA. The docking parameters used were: maximum number of energy evolutions: 250,000; GA population size: 150; and the number of GA runs; 30 [26]. The conformer with the lowest binding energy was used for further studies and it was in sync with the experimental results obtained by Fluorescence spectroscopy.

### Molecular dynamics

Molecular dynamics (MD) Simulation for 10 ns was performed at 300K and 1 bar pressure for the energetically most stable conformer by using GROMACS V 4.6.3 software. The free energy obtained from fluorescence and the best conformer from docking results was used for the MD simulations. The topology parameters of HSA were created by using Dundee PRODRG2.5 server (beta). The complex structure was immersed in a box of 7.335 × 6.135 × 8.119 nm^3^dimension with extended simple point charge (SPC) water molecules. Sodium counter ions were added to maintain electro-neutrality. All the parameters were same as our previously reported work [27]. Simulation and analysis were performed on Linux cluster with 36 nodes (dual Xeon processor) at the Bioinformatics facility, University of Hyderabad.

## Results

### Fluorescence spectroscopy

Since Ashwagandha is used as Ayurveda medicine, however, the active principles of these phytochemicals withanolide A, withanolide B, withanoside IV and withanoside V were not been understood well related to food supplements. Since, HSA is an important protein to transport the small molecules to the target places, thus, we have focused our study to understand the binding mechanism of these isolated molecules with HSA. The intrinsic fluorescence in HSA is because of the presence of a single tryptophan residue at the position 214, also there are 18 tyrosine and 33 phenylalanine residues. By continuous titration of phytochemicals withanolide A, withanolide B, withanoside IV and withanoside V to HSA, the fluorescence of protein was quenched. These results inferthat the intrinsic fluorescence of HSA is quenched gradually with increasing concentrations of withanolides and withanosides derivatives because of the change in microenvironment around Trp 214 residue. Its inner filter affect can be corrected by using the following equation,

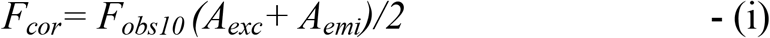

Where F_cor_ and F_obs_ are corrected and observed fluorescence intensity respectively. A_ext_ and A_emi_ are absorbance at fluorescence excitation (285 nm) and emission (360 nm) wavelengths [28]. To validate whether the complex is undergoing dynamic or static quenching, bimolecular quenching constant was calculated from the slope of F_O_/F vs Q and it was found to be linear, which indicates that it is undergoing static quenching. The fluorescence data was analyzed using the stern-volmer equation

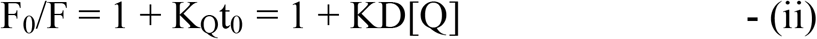

where F_0_ and F are fluorescence intensities, in the absence and presence of the quencher. Q is quencher concentration and K_D_ is the stern-volmer quenching constant which can be written as K_D_ =Kqt_0_; t_0_ is the lifetime of the fluorophore for HSA (5.6ns). Since the interaction between withanolides and withanosides, and HSA is by static mode of binding, the modified stern-volmer regression curve is used to determine the binding constant (Ks) and the number of binding sites (n), where n is the slope and Ks is the binding constant.

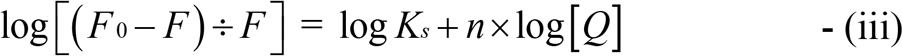

The binding constants were calculated to be K_Withanolide A_ = 3.04 ±0.05 × 10^4^M^−1^, K_Withanolide B_= 7.59±0.05 × 10^4^M^−1^, K_Withanoside IV_ = 6.74±0.03 × 10^4^M^−1^and K_Withanoside V_ = 5.33±0.05 × 10^4^M^−1^respectively as shown in (Fig 1A-D). This data indicates that withnolide and withanoside derivatives are strongly binding to the HSA. Hence, our results are in agreement with previously published phytochemicals were strongly associated with HSA [29–31]. Also, the obtained binding constants are in concurrence with the food and drug administration (FDA) which indicates that phytochemicals used here withanolide A, withanolide B, withanoside IV and withanoside V could be potential therapeutic molecules.

**Fig 1.**
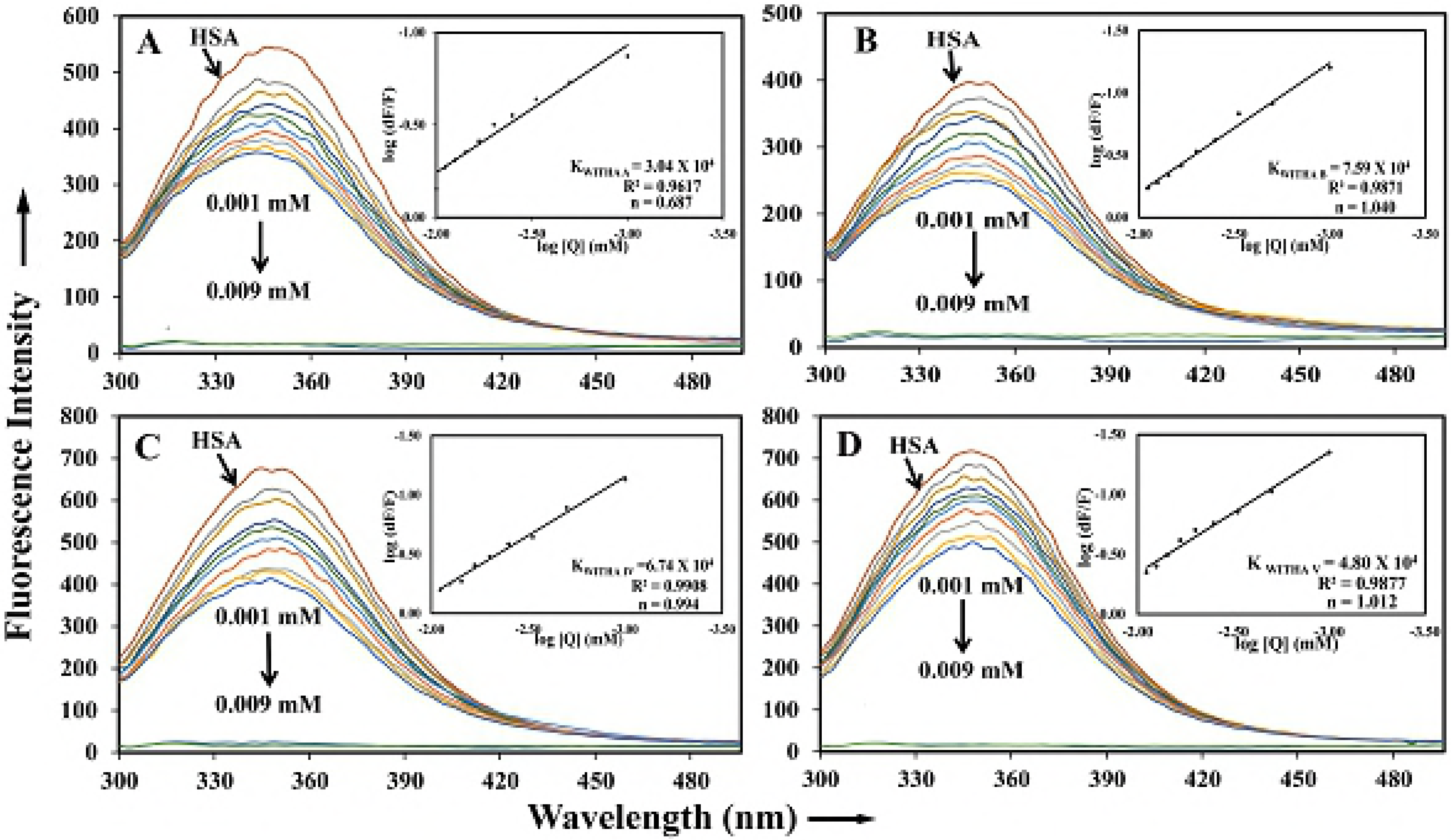
Fluorescence spectroscopic studies of HSA with withanolide and withanoside molecules, indicating the interaction of the drug with plasma protein. The association constant (**K_S_**) and free energy change along with stern-volmer plots showing fluorescence quenching constant (kq) and plot of Fo/F against [Q] at λex = 285 nm and λem = 360 nm for (A) withanolide A, (B) withanolide B, (C) withanoside IV and (D) withanoside V.

The standard free energy change is calculated by using the following equation.

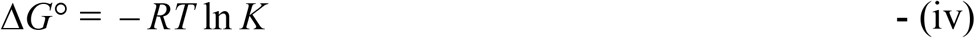

Where ΔG is the free energy change, R is the gas constant at room temp and K is binding constant calculated from fluorescence data. The free energy changes upon binding of withanolides and withanosides with HSA was −5.61 Kcal M^−1^, −6.63 Kcal M^−1^, −6.56 Kcal M^−1^and −6.36 Kcal M^−1^respectively at 25 °C. This indicates that the interaction between the drugs with serum albumin is mainly hydrophobic interactions. The computationally derived free energy change was found to be totally in sync with the experimental data.

### Displacement studies

Using site specific markers we can understand the exact binding of ligand molecules to the specific domains of HSA. Hence, there are different site specific markers like lidocaine for domain I, phenylbutazone for domain II and ibuprofen for domain III [22] and using these markers the fluorescence was performed to analyze the specific binding domain of HSA on interaction with withanoside and withanolide derivatives [28,32]. Because of the structural and molecular similarity of four derivatives, all of them showed fluorescence emission quenching by displacing phenylbutazone, i.e. they are binding on domain II of HSA with a binding constant were K_WithanolideA+pb_ = 2.57±0.05 × 10^4^M^−1^, K_WithanolideB+pb_ = 6.84±0.05 X10^4^M^−1^, K_WithanosideIV+pb_ = 1.89±0.05 × 10^4^M^−1^and K_WithanosideV+pb_= 4.80±0.03 × 10^4^M^−1^. The free energy changes for different were −6.09 Kcal M^−1^, −6.56 Kcal M^−1^, −7.17 Kcal M^−1^and −6.42 Kcal M^−1^respectively (Fig 2 A-D). We also performed with other site specific markers (i.e lidocaine, ibuprofen), however, they were not displaced by the withnolide compounds. These results indicate that the drug molecules are specifically binding to Sudlow’s drug binding site I. Our Experimental results are in congruence with the computational data as also illustrated in our previous reports [33].

**Fig 2.**
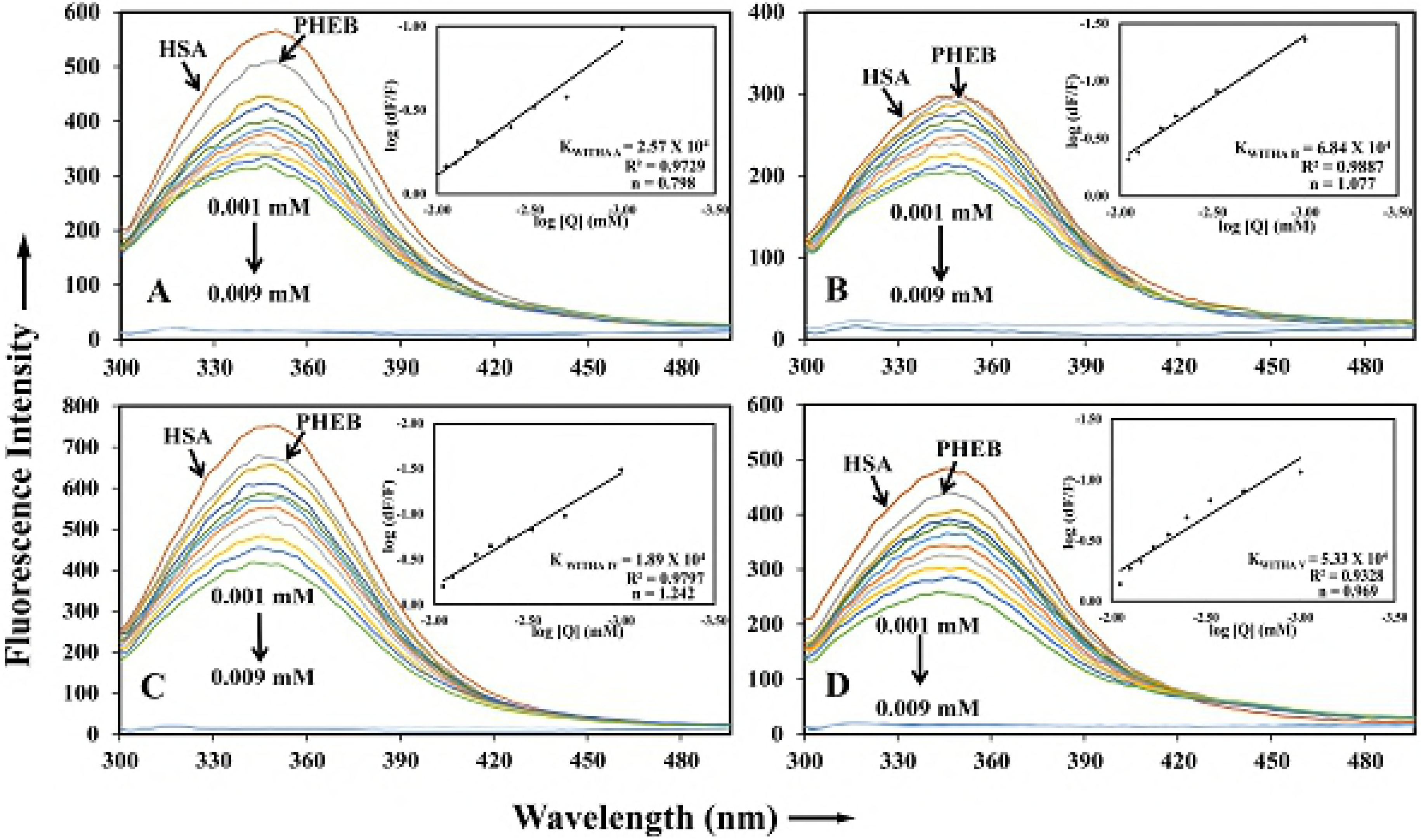
Site displacement studies using site-specific markers; Phenylbutazone was used as marker for HSA domain IIA (Sudlow site I) Fluorescence spectroscopic studies performed using HSA and Phenylbutazone at equal concentrations (1μM) and drugs with increasing concentrations (1μM ~ 9μM).

### Circular dichroism

Circular dichroism is a fundamental technique to study any secondary structural change upon interaction of protein and ligand. HSA showed a dip at 208 and 222nm in the far UV region which are mainly originated from helical structure [20]. The secondary structure of HSA comprises of 58% α-helix, 20% β-sheet (parallel and anti-parallel) and 22% random coils, but upon titration with withanolide and withanoside molecules there was partial unfolding of HSA protein and change in the dip in 208 and 222nm as shown in (Fig 3 A-D).

**Fig 3.**
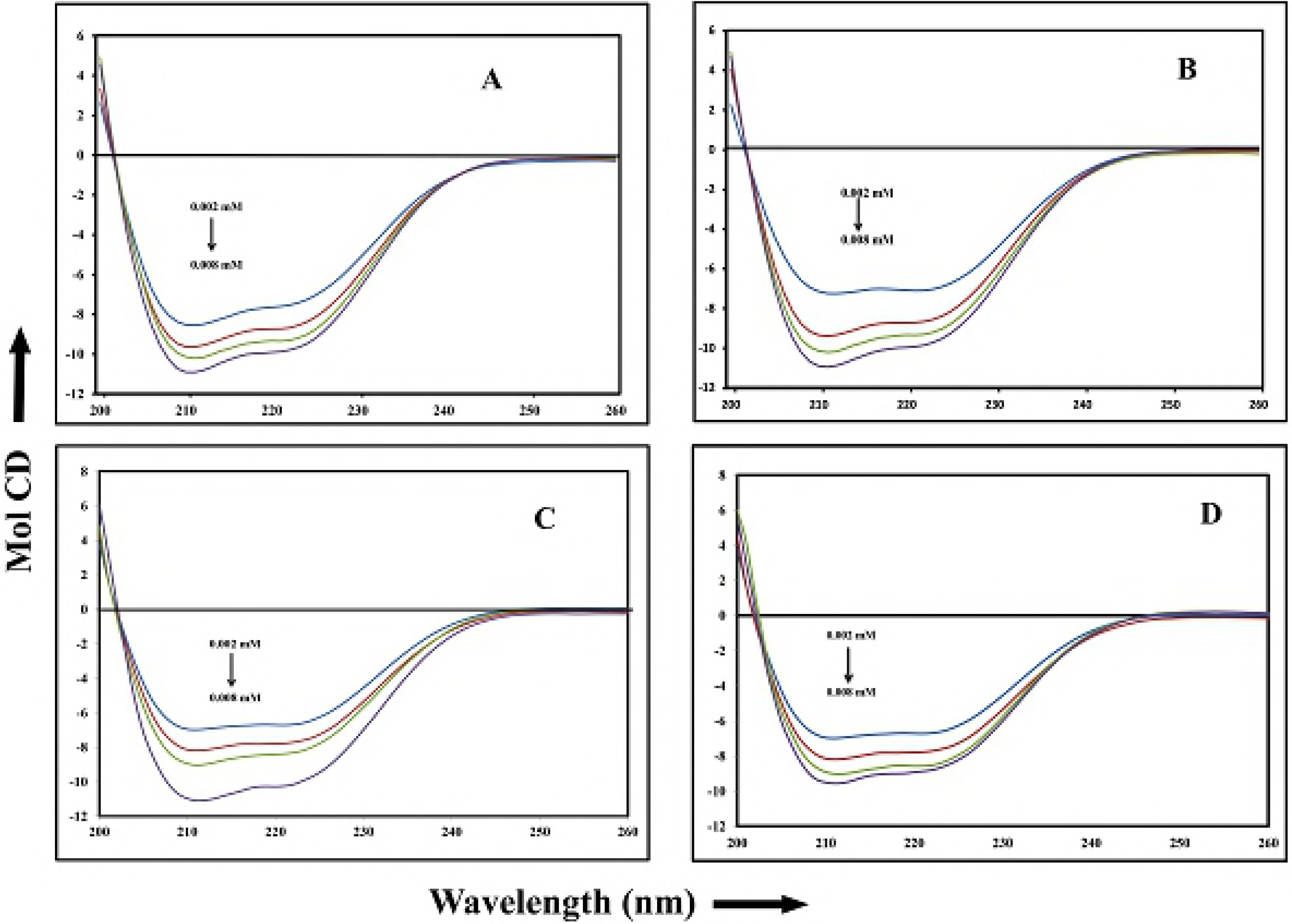
Circular Dichroism studies of the free HSA and HSA-drug complexes. The free HSA and HSA-drug complexes in aqueous solution with a protein concentration fixed at 1μM and with increasing drug concentrations at 2,4, and 6 μM. (A) withanolide a, (B) withanolide b, (C) withanoside IV and (D) withanoside V.

Conformational analysis was done using CDNN software, shows the percentage of α-helix increasing up to 69.30±2.5, 70.54±2.5, 66.8±2.5 and 61.5±2.3 upon binding with withanolide A, withanolide B, withanoside IV and withanoside V respectively, and simultaneously there is decrease in the percentage of β-sheet and random coil (S Table 1). In general most of the ligand molecules binding to HSA [34,35]. Similar studies were done for various molecules and revealed that upon binding of ligands, there is change in the secondary structure of HSA [36].

### Topological observations from atomic force microscopy

To corroborate topological changes in surface of free HSA and HSA upon addition of drug derivatives, atomic force microscopy (AFM) was used. The results explain that upon incubation with drug molecules, there is a significant increase in the size of the complex as shown in (Fig 4 A-J). But as the molecular weight of withanoside IV and withanoside V is larger than that of withanolide A and withanolide B, the complex formed by interaction with withanoside IV and withanoside V are larger. The unliganded HSA is showing a small size comparatively with bound HSA with withanaloides, and the results corroborate with the previous studies [37]. The complex in the presence of withanolide (A, B), and withanoside (IV, V) were showing remarkable increase in size i.e. growing around to become 70nm, 130nm, 190 nm and 300 nm (S1 Fig). These results indicate that the above molecules are formed complexes with HSA. Our results deciphered that it could be the hydrophobic contacts play a major role while binding of withanolides and withanosides compounds with HSA complexation [38] which is in an agreement with the free energy calculations. These experiments were performed in triplicates and similar results were reproduced always.

**Fig 4.**
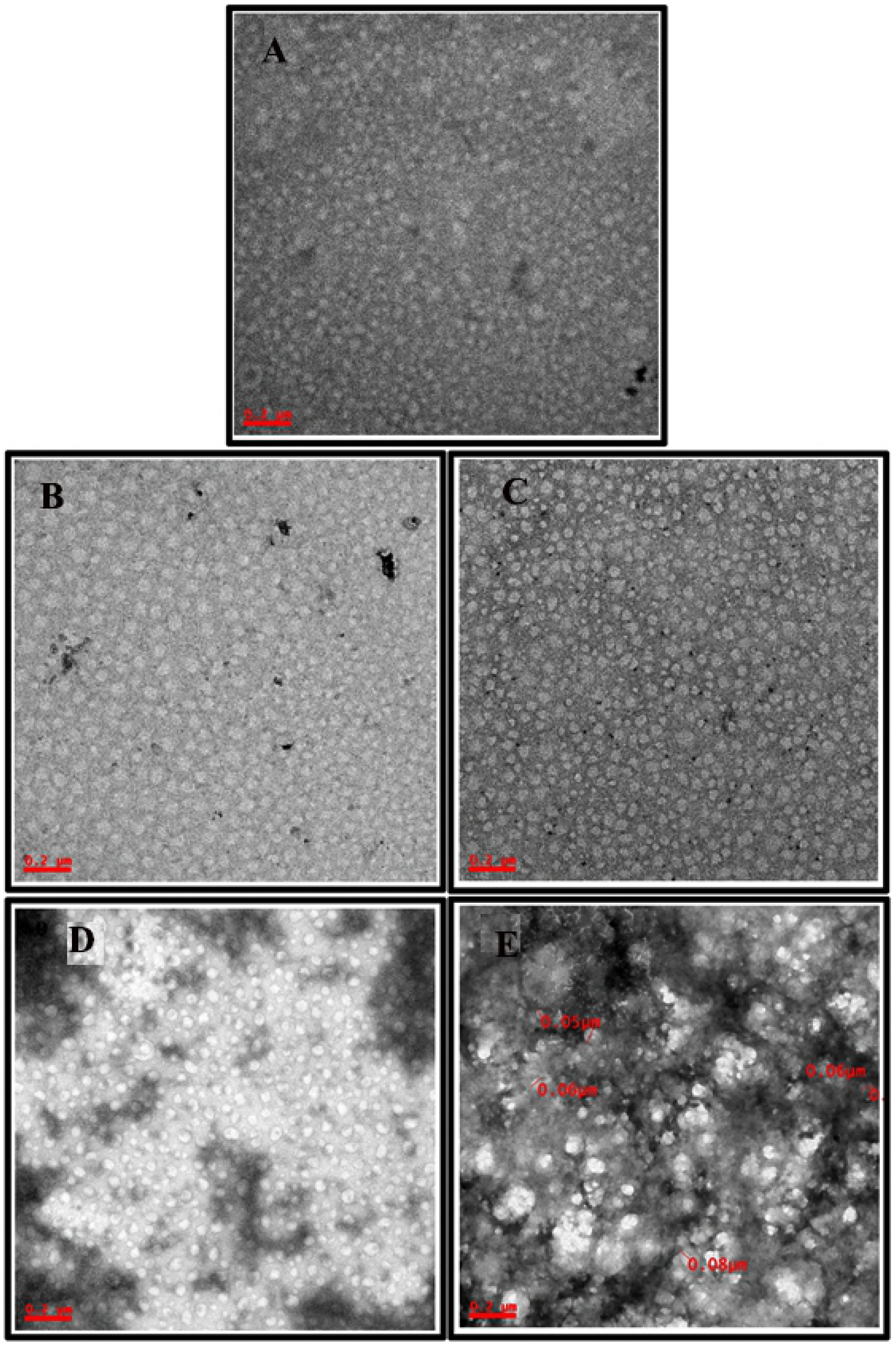
Atomic Force Microscopic (AFM) studies to visualize alteration in HSA molecule topology in presence of withanolide and withanoside derivatives at 10 μM resolution. (A) ONLY has (B) HSA+withanolide A (C) HSA+withanolide B (D) HSA+withanolide IV and (E) HSA+withanolide V.

### Transmission emission microscopy

TEM was used to visualize the structural and topographical change in HSA and ligated HSA upon incubation with various drug derivatives [39]. At resolution of 0.2μM the size of unligated HSA molecule were found to be of size 0.04±10μM, whereas in the presence of withanolide A and B, and withanoside IV and V, it increased to 0.06±10μM, 0.06±10μM, and 0.07±10μM, 0.08±10μM respectively (Fig 5 A-E), which can interprets the complexes formed by the interaction of HSA with these derivatives. Thus, it can be derived that interaction among the protein and withanolide A, withanolide B, withanoside IV and withanoside V are taking place and these results are in harmony with the results obtained from other techniques.

**Fig 5.**
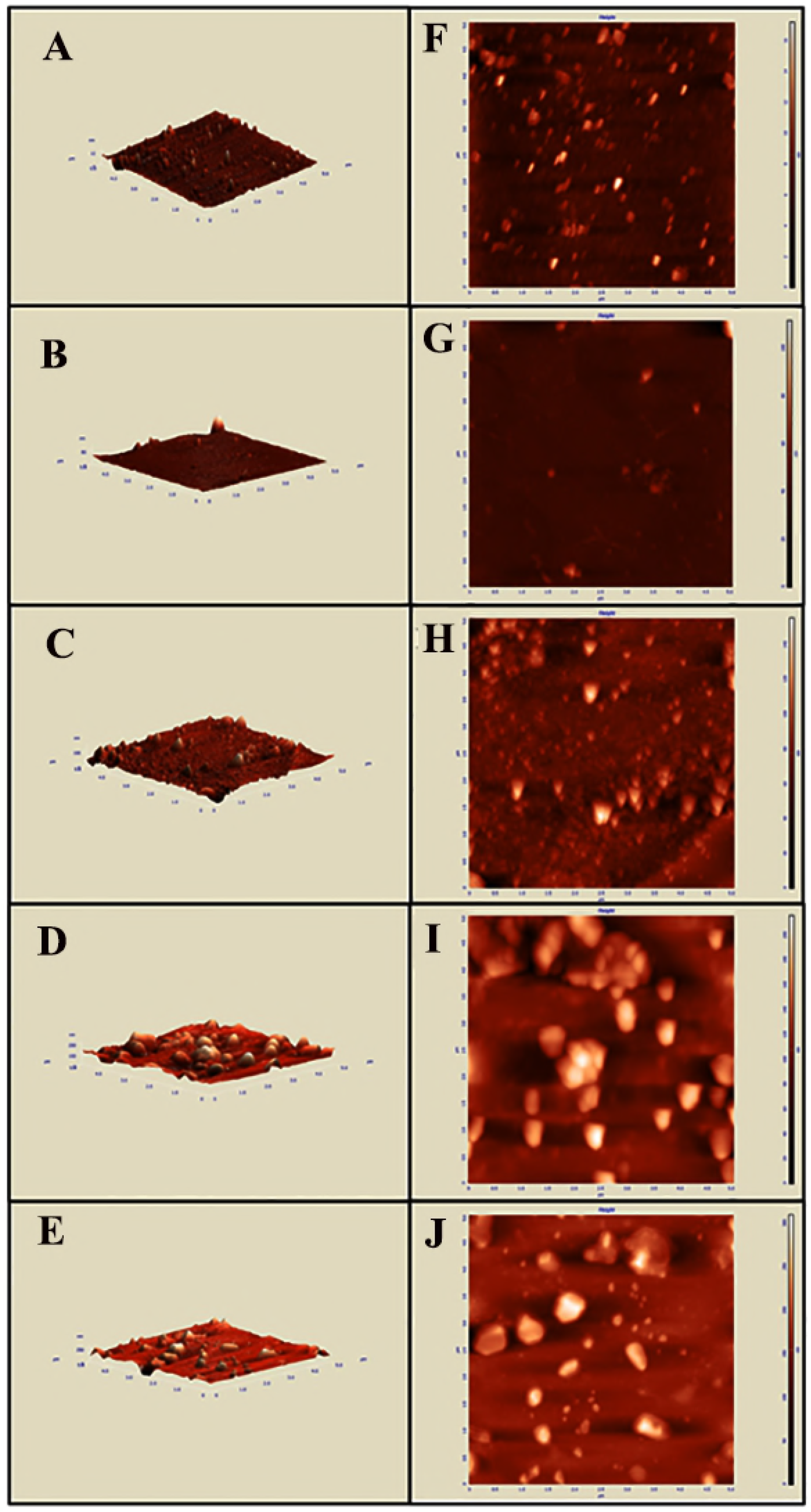
Transmission Electron Microscopic (TEM) studies to visualize alteration in HSA molecule topology in presence of withanolide and withanoside derivatives at 200 nM resolution. (A) ONLY HSA (B) HSA+withanolide A, (C) HSA+withanolide B, (D) HSA+withanolide IV and (E) HSA+withanolide V.

### Molecular docking

The displacement studies by the use of site specific markers indicated that the withanolide and withanoside derivatives were binding at Sudlow’s drug binding site I of HSA. Autodock 4.2.3 software was used further to confirm the precise binding site and residues on HSA upon binding of these derivatives. Since the binding location of the protein is of utmost importance to study the biological activity of the drug and it also plays a major role in pharmacokinetics and pharmacodynamics of the drug. Crystal structure of HSA was procured from Protein Data Bank (PDB Code: 1A06). Withanolide A is binding to HSA by hydrogen bond formation by interacting with Glu208 in the hydrophobic cavity of subdomain IIA, and withanolide B is interacting with His338, withanoside IV is forming a hydrogen bond at Arg197 and Lys205, and withanoside V is interacting with residues of Tyr341 and Tyr334 and Leu305; with binding constants of K_Withanolide A_ = 4.93 × 10^4^M^−1^, K_Withanolide B_ = 7.42 × 10^4^M^−1^, K_Withanoside IV_ = 2.50 × 10^4^M^−1^and K_Withanoside V_ = 2.49 × 10^4^M^−1^and free energy were be −6.40 Kcal M^−1^, −6.64 Kcal M^−1^, −3.27 Kcal M^−1^and −3.26 Kcal M^−1^respectively at 25 °C respectively. The results were shown in (Fig 6A-L), and are generated by using the Pymol software, and Ligplot is used to illustrate the two-dimensional interaction by hydrogen bond formation and hydrophobic interactions. The Binding constant values calculated computationally were in accordance with the values obtained experimentally [32,40].

**Fig 6.**
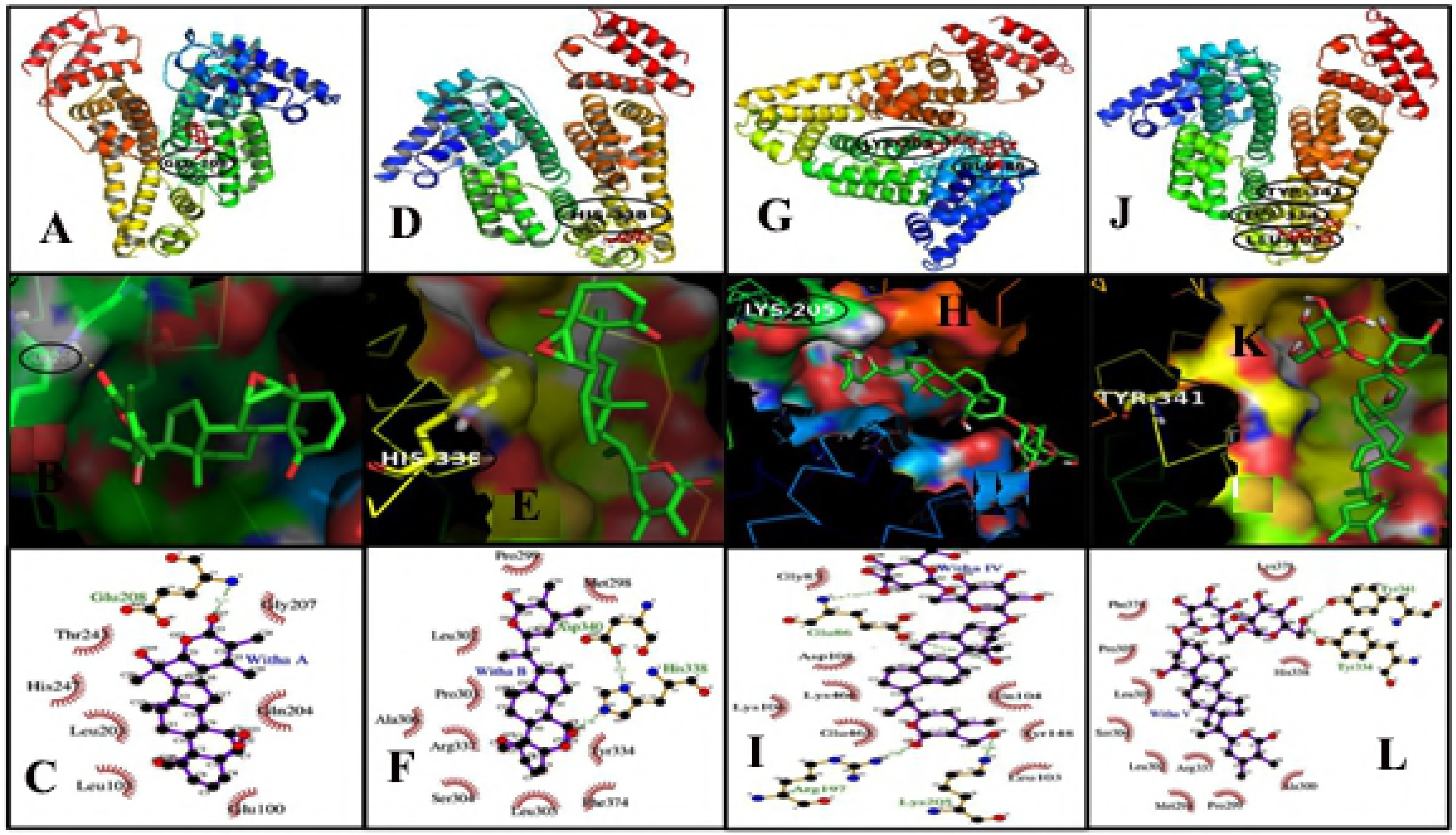
Molecular docking studies between HSA and withanolide A, withanolide B, withanoside IV and withanoside V showed that the minimum binding energy conformer is very close to the experimentally determined values. (A-L) Different views of the derivatives docked in the binding pocket HSA using Autodock 4.2. Overview of cartoon model of withanolide A, withanolide B, withanoside IV and withanoside V binding to HSA. Pymol is used to generate the images. The cavity of hydrophobic and hydrophilic amino acid residues surrounding the probe. Ligplot is used to show the hydrophobic interactions of HSA with drug derivatives.

### Molecular dynamics

Molecular simulation studies compute the behaviour of a system as a function of time. It has emerged as a powerful tool for understanding the interaction between protein and ligands to predict how conformational changes occur to achieve the lowest free energy conformer. Usually, RMSD (root mean square deviation), RMSF (Root mean square fluctuation) and Rg (radius of gyration) are used to know the change in the microenvironment, atomic fluctuation, rigidity and stability of the HSA-ligand complexes in comparison to HSA alone [41,42]. Hence, in our study we have measured the same parameters for HSA with withanolide and withanoside complexes to understand the interaction mechanism. Energetically the most stable complex of docking was taken and dynamics was studied for 10ns.

Generally, Rg is used for measuring the compactness of structure, hence Rg of withanolides and withanosides complexes showed stability throughout the 10ns after initial rigidity at 2ns. The Rg value of unliganded HSA is, 2.65±0.35nm, the complex HSA-withanolide A, HSA-withanolide B, HSA-withanoside IV and HSA-withanoside V showed fluctuations in between 2.63±0.05nm, 2.61±0.02nm, 2.65±0.03nm, and 2.65±0.02nm, respectively showing the stability of the complexes. These results indicate the conformational and stability changes in the secondary structure of HSA, which is totally in congruence with the results obtained from circular dichroism.

To access the stability of the system RMSD of the complex atoms was analysed as a function of time for MD trajectory. The RMSD values of atoms of protein backbone (C-Cα-N) were calculated for only HSA and HSA-ligand complex. RMSD of HSA-withanolide A, HSA-withanolide B, HSA-withanoside IV, and HSA-withanoside V showed stability at 3ns and remained constant throughout 10ns. From 0-10ns trajectory data the RMSD value of unliganded HSA is 0.4±0.03nm while for HSA-ligand complexes fluctuations were in between 0.3±0.03nm, while withanolide B had initial fluctuations which later stabilized at 6.5 ns from 0.35±0.03nm as shown in (Fig 7A-L). It can be concluded that the complexes remain stable with no major change from the initial docked conformer owing to the stability of the ligated HSA.

**Fig 7.**
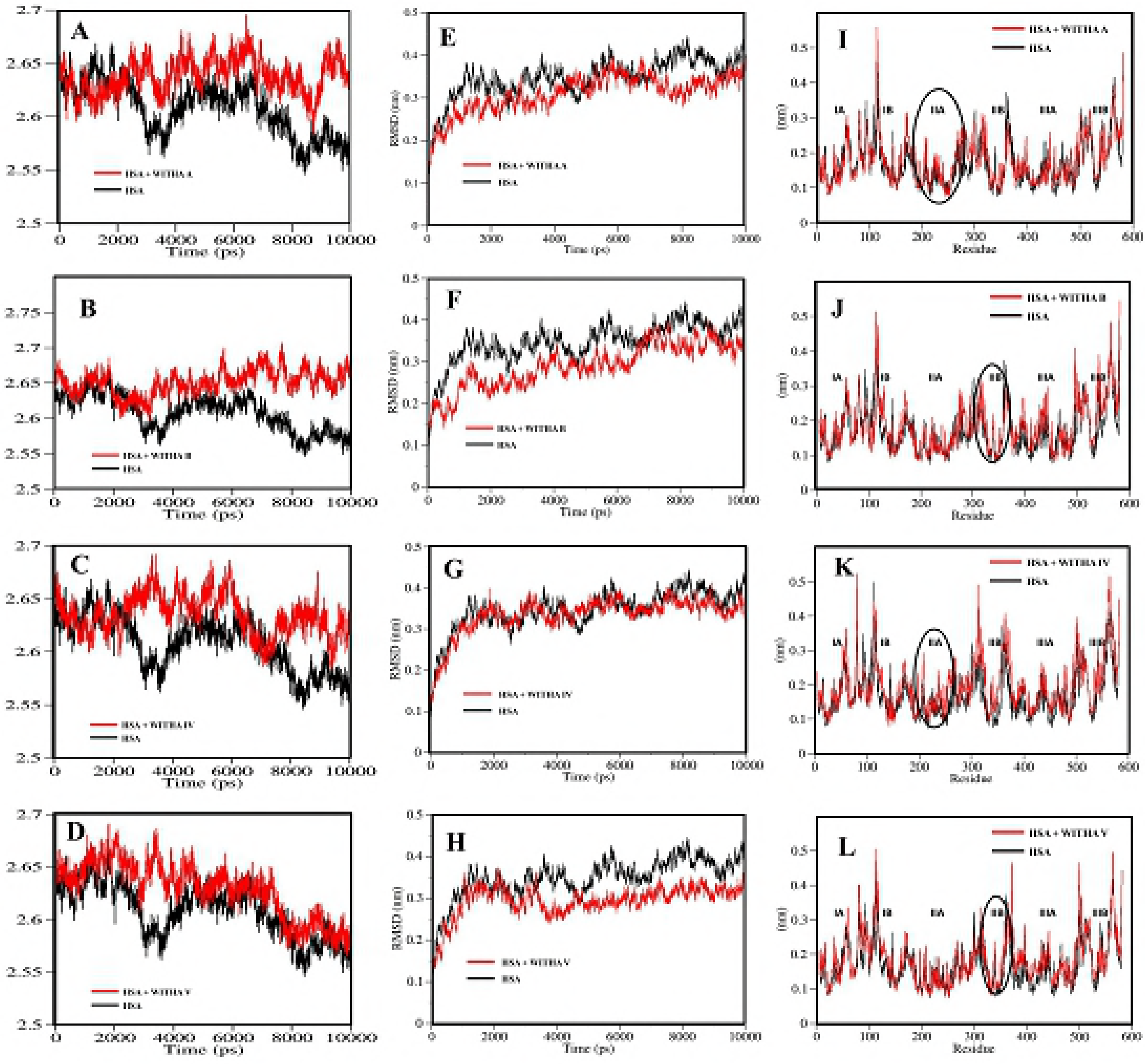
(A-D)-Time evolution of the radius of gyration (Rg) during 10 ns of MD simulation of unliganded HSA and HSA-drug derivatives complexes. (E-H)-Plot of RMSD values for unliganded HSA and HSA-drug derivatives complexes. (I-L)-Comparison of the RMSF of Calcium atoms along the sequence derived from the 10 ns simulations

Rigidity of peptide structure and thermal vibrations were measured by RMSF values. Local protein mobility was seen by analysing the time averaged RMSF values of only HSA and the HSA-drug complexes. As these derivates of withanolide A, withanolide B, withanoside IV and withanoside V binds to HSA, there is increased rigidity and flexibility in different sub domains of HSA which is plotted against the residue number. The microenvironment around the binding site showed strong overlapping of binding region of the HSA-withanolide A, HSA-withanolide B, HSA-withanoside IV and HSA-withanoside V and are in between 200-250 residues which are a part of hydrophobic cavity of HSA and there is more rigidity around those amino acids indicating strong interaction at binding site 1(S Fig 2). Hence, MD data is an indication that the withanolide A, withanolide B, withanoside IV and withanoside V derivatives are binding with HSA with stable conformations and hence this study can be extend to study the therapeutic role in biological samples.

## Conclusion

The goal of this study was to elucidate the effect of interaction of phytocompounds, i.e. derivatives of *Withania somnifera* (Ashwagandha) namely withanolide A, withanolide B, withanoside IV and withanoside V, with carrier plasma protein HSA. Quenching of fluorescence emission of HSA showed the formation of HSA-ligand complex formation with binding constant of K_Withanolide A_ = 3.04±0.05 × 10^4^M^−1^, K_Withanolide B_ = 7.59±0.05 × 10^4^M^−1^, K_Withanoside IV_ = 6.74±0.03 × 10^4^M^−1^and K_Withanoside V_ = 5.33±0.05 × 10^4^M^−1^. These binding constants fall under the range of 10^3^ - 10^6^M^−1^, which is in the range of known FDA approved drugs. Circular Dichroism also shows partial unfolding of the protein upon interaction of these molecules with HSA and further it was confirmed by AFM and TEM on basis of morphological and topological changes in the protein-ligand complexes. Our experimental results illustrated strong binding between these derivatives with HSA and is in corroboration with the *in silico* data of molecular docking and MD simulations. Owing to similar molecular and structural formula all the derivatives of *Withania somnifera,* were seen to displace phenylbutazone and bind on Sudlow’s site I. Using MD simulation the stability of only HSA and HSA-drug complexes were qualitatively compared for 10 ns. The study undertaken in our lab will be helpful in further understanding the pharmacokinetics and pharmacodynamics of these compounds and it provides a base to further exploit the far-reaching pharmaceutical potential of these steroidal derivatives.

## AUTHOR CONTRIBUTIONS

Designed the experiments: SD, RS, SD and AS performed the experiments (compound binding analysis), KM did Molecular simulations. Manuscript preparation was done by SD and RS. All authors have read and approved the final manuscript.

## ACKNOWLEDGEMENTS

This work was supported by Science and Engineering Research Board (SB/EMEQ-064/2014 dated 14-07-2015 and DST-FIST) and UGC-SAP. We thank BIF, for Bioinformatics facilities. SD and MK acknowledges CSIR and UGC-Kothari for fellowship.

## CONFLICT OF INTEREST

The authors have declared that there is no conflict of interest.

## References

1. Glotter E. Withanolides and related ergostane-type steroids. Nat Prod Rep. 1991 Aug 8; 4:415–40. PMID: 1787922

2. Dhar N, Razdan S, Rana S, Bhat WW, Vishwakarma R, et al. A Decade of Molecular Understanding of Withanolide Biosynthesis and In vitro Studies in Withania somnifera (L.) Dunal: Prospects and Perspectives for Pathway Engineering. Front Plant Sci. 2015 Nov 27; 6:1031. doi: 10.3389/fpls.2015.01031 PMID: 26640469

3. Mirjalili MH, Moyano E, Bonfill M, Cusido RM, Palazon J. Steroidal lactones from Withania somnifera, an ancient plant for novel medicine. Molecules. 2009 Jul 3; 14:2373–93. doi: 10.3390/molecules14072373 PMID: 19633611

4. Verma S. Therapeutic uses of Withania somnifera (Ashwagandha) with a note on withanolides an its pharmacological actions. Asian J Pharm Clinical Res. 2011 Jul 4:1

5. Mishra LC, Singh BB, Dagenais S. Scientific basis for the therapeutic use of Withania somnifera (ashwagandha): a review. Altern Med Rev 2000 Aug 5; 4: 334–346. PMID: 10956379

6. Chuang VT, Otagiri M. Stereoselective binding of human serum albumin. Chirality. 2006 Feb 18; 3:159–66. DOI: 10.1002/chir.20237 PMID: 16432913.

7. Peters T, Jr. Serum albumin. Adv Protein Chem. 1985; 37:161–245. PMID: 3904348

8. Carter DC, He XM, Munson SH, Twigg PD, Gernert KM, et al. Three-dimensional structure of human serum albumin. Science. 1989 Jun 9; 244:1195–8. PMID: 2727704

9. Peters TJ. All About Albumin: Academic Press. Dec 8; 432. ISBN: 9780080527048

10. Sudlow G, Birkett DJ, Wade DN (1976) Further characterization of specific drug binding sites on human serum albumin. Mol Pharmacol. 1976 Nov 12; 6:1052–61. PMID: 1004490

11. Zsila F. Subdomain IB Is the Third Major Drug Binding Region of Human Serum Albumin: Toward the Three-Sites Model. Mol Pharm. 2013 May 6; 10:1668–82. doi: 10.1021/mp400027q PMID: 23473402

12. Curry S, Brick P, Franks NP. Fatty acid binding to human serum albumin: new insights from crystallographic studies. Biochim Biophys Acta. 1999 Nov 23; 1441:131–40 PMID: 10570241

13. Varshney A, Sen P, Ahmad E, Rehan M, Subbarao N, et al. Ligand binding strategies of human serum albumin: how can the cargo be utilized? Chirality. 2010 Jan 22; 1:77–87. doi: 10.1002/chir.20709 PMID: 19319989

14. Gokara M, Sudhamalla B, Amooru DG, Subramanyam R. Molecular interaction studies of trimethoxy flavone with human serum albumin. PLoS One. 2010 Jan 21; 5:e8834. doi: 10.1371/journal.pone.0008834 PMID: 20098677

15. Gokara M, Malavath T, Kalangi SK, Reddana P, Subramanyam R. Unraveling the binding mechanism of asiatic acid with human serum albumin and its biological implications. J Biomol Struct Dyn. 2014; 32:1290–302. doi: 10.1080/07391102.2013.817953 PMID: 23844909

16. Kallubai M, Rachamallu A, Yeggoni DP, Subramanyam R. Comparative binding mechanism of lupeol compounds with plasma proteins and its pharmacological importance. Mol Biosyst. 2015 Apr 11; 4:1172–83. doi: 10.1039/c4mb00635f. PMID: 25710711

17. Sudhamalla B, Gokara M, Ahalawat N, Amooru DG, Subramanyam R. Molecular dynamics simulation and binding studies of beta-sitosterol with human serum albumin and its biological relevance. J Phys Chem B. 2010 Jul 15; 114:9054–62. doi: 10.1021/jp102730p PMID: 20565066

18. Gokara M, Narayana VV, Sadarangani V, Chowdhury SR, Varkala S, et al. Unravelling the binding mechanism and protein stability of human serum albumin while interacting with nefopam analogues: a biophysical and insilico approach. J Biomol Struct Dyn. 2017 Aug 16; 35:2280–2292. doi: 10.1080/07391102.2016.1216895 PMID: 27453381

19. Yeggoni DP, Rachamallu A, Subramanyam R. Protein stability, conformational change and binding mechanism of human serum albumin upon binding of embelin and its role in disease control. J Photochem Photobiol B. 2016 Apr 18; 160:248–59. doi: 10.1016/j.jphotobiol.2016.04.012 PMID: 27130964

20. Subramanyam R, Gollapudi A, Bonigala P, Chinnaboina M, Amooru DG. Betulinic acid binding to human serum albumin: a study of protein conformation and binding affinity. J Photochem Photobiol B. 2009 Jan 9; 94:8–12 doi: 10.1016/j.jphotobiol.2008.09.002 PMID: 18945624

21. Yamasaki K, Rahman MH, Tsutsumi Y, Maruyama T, Ahmed S, et al. Circular dichroism simulation shows a site-II-to-site-I displacement of human serum albumin-bound diclofenac by ibuprofen. AAPS PharmSciTech. 2000 May 14; 1:E12 doi: 10.1208/pt010212 PMID: 14727845

22. Ghuman J, Zunszain PA, Petitpas I, Bhattacharya AA, Otagiri M, et al. Structural basis of the drug-binding specificity of human serum albumin. J Mol Biol. 2005 Oct 14; 353:38–52 DOI: 10.1016/j.jmb.2005.07.075 PMID: 16169013

23. Hein KL, Kragh-Hansen U, Morth JP, Jeppesen MD, Otzen D, et al. Crystallographic analysis reveals a unique lidocaine binding site on human serum albumin. J Struct Biol. 2010 Sep; 171:353–60 doi: 10.1016/j.jsb.2010.03.014 PMID: 20347991

24. Yeggoni DP, Manidhar DM, Suresh Reddy C, Subramanyam R. Investigation of binding mechanism of novel 8-substituted coumarin derivatives with human serum albumin and alpha-1-glycoprotein. J Biomol Struct Dyn. 2016 Sep; 34:2023–36 doi: 10.1080/07391102.2015.1104264 PMID: 26440860

25. Yeggoni DP, Rachamallu A, Subramanyam R. A comparative binding mechanism between human serum albumin and [small alpha]-1-acid glycoprotein with corilagin: biophysical and computational approach. RSC Advances. Apr 6; 6: 40225–40237 doi: 10.1039/C6RA06837E

26. Yeggoni DP, Rachamallu A, Kallubai M, Subramanyam R. Cytotoxicity and comparative binding mechanism of piperine with human serum albumin and alpha-1-acid glycoprotein. J Biomol Struct Dyn. 2015 Aug 20; 33:1336–51. doi: 10.1080/07391102.2014.947326 PMID: 25054206

27. Yeggoni DP, Subramanyam R. Binding studies of L-3,4-dihydroxyphenylalanine with human serum albumin. Mol Biosyst. 2014 Dec; 10:3101–10. doi: 10.1039/c4mb00408f PMID: 25209359

28. Yadav SA, Yeggoni DP, Devadasu E, Subramanyam R. Molecular binding mechanism of 5-hydroxy-1-methylpiperidin-2-one with human serum albumin. J Biomol Struct Dyn. 2018 Mar 22; 36:810–817. doi: 10.1080/07391102.2017 PMID: 28278025

29. Agudelo D, Bourassa P, Bruneau J, Berube G, Asselin E, et al. Probing the binding sites of antibiotic drugs doxorubicin and N-(trifluoroacetyl) doxorubicin with human and bovine serum albumins. PLoS One. 2012 Aug 24; 7:e43814. doi: 10.1371/journal.pone.0043814 PMID: 22937101

30. Naveenraj S, Anandan S. Binding of serum albumins with bioactive substances – Nanoparticles to drugs. J Photochem Photobiol Photochem Rev. 2013 Mar; 14: 53–71 doi.org/10.1016/j.jphotochemrev.2012.09.001

31. Gokara M, Kimavath GB, Podile AR, Subramanyam R. Differential interactions and structural stability of chitosan oligomers with human serum albumin and alpha-1-glycoprotein. J Biomol Struct Dyn. 2015 Dec 20; 33: 196–210 doi: 10.1080/07391102.2013.868321 PMID: 24359035

32. Yeggoni DP, Kuehne C, Rachamallu A, Subramanyam R. Elucidating the binding interaction of andrographolide with the plasma proteins: biophysical and computational approach. RSC Adv. Dec 9; 7: 5002–5012. doi: 10.1039/C6RA25671F

33. Nerusu A, Srinivasa Reddy P, Ramachary D, Subramanyam R. Unraveling the Stability of Plasma Proteins upon Interaction of Synthesized Androstenedione and Its Derivatives—A Biophysical and Computational Approach. ACS Omega 2017 Oct 9; 2: 6514–6524 doi: 10.1021/acsomega.7b00577.

34. Yuan L, Liu M, Liu G, Li D, Wang Z, et al. Competitive binding of (–)-epigallocatechin-3-gallate and 5-fluorouracil to human serum albumin: A fluorescence and circular dichroism study. Spectrochimica Acta Part A: Molecular and Biomolecular Spectroscopy. Spectrochim Acta A Mol Biomol Spectrosc. 2017 Feb 15; 173: 584–592 doi: 10.1016/j.saa.2016.10.023 PMID: 27776313

35. Subramanyam R, Goud M, Sudhamalla B, Reddeem E, Gollapudi A, et al. Novel binding studies of human serum albumin with trans-feruloyl maslinic acid. J Photochem Photobiol B. 2009 May 4; 95:81–8 doi: 10.1016/j.jphotobiol.2009.01.002 PMID: 19230701

36. Neelam S, Gokara M, Sudhamalla B, Amooru DG, Subramanyam R. Interaction studies of coumaroyltyramine with human serum albumin and its biological importance. J Phys Chem B. 2010 Mar 4; 114:3005–12. doi: 10.1021/jp910156k PMID: 20136105

37. Kallubai M, Reddy SP, Dubey S, Ramachary DB, Subramanyam R. Spectroscopic evaluation of synthesized 5β-dihydrocortisol and 5β-dihydrocortisol acetate binding mechanism with human serum albumin and their role in anticancer activity. J Biomol Struct Dyn. 2018 Feb; 15: 1–18. doi: 10.1080/07391102.2018.1433554 PMID: 29375009

38. Chanphai P, Froehlich E, Mandeville JS, Tajmir-Riahi HA. Protein conjugation with PAMAM nanoparticles: Microscopic and thermodynamic analysis. Colloids Surf B Biointerfaces. 2017 Feb 1; 150:168–174 doi: 10.1016/j.colsurfb.2016.11.037 PMID: 27914253

39. Huang J, Xie J, Chen K, Bu L, Lee S, et al. HSA coated MnO nanoparticles with prominent MRI contrast for tumor imaging. Chem Commun (Camb). 2010 Sep 28; 46:6684–6. doi: 10.1039/c0cc01041c PMID: 20730157

40. Li S, He J, Huang Y, Wang Q, Yang H, et al. Interactions of cucurbit[6,7]urils with human serum albumin and their effects on zaltoprofen transportation. RSC Adv 2016 Sep 3; 6: 85811–85819 doi: 10.1039/C6RA17508B.

41. Rapaport DC, Blumberg, Robin L., McKay, Susan R., Christian, Wolfgang. The Art of Molecular Dynamics Simulation. Comput Phys Sep; 10:456 doi: 10.1063/1.4822471.

42. Radibratovic M, Minic S, Stanic-Vucinic D, Nikolic M, Milcic M, et al. Stabilization of Human Serum Albumin by the Binding of Phycocyanobilin, a Bioactive Chromophore of Blue-Green Alga Spirulina: Molecular Dynamics and Experimental Study. PLoS One. 2016 Dec 13; 11:e0167973. doi: 10.1371/journal.pone.0167973 PMID: 27959940s

